# HMeta-d: Hierarchical Bayesian estimation of metacognitive efficiency from confidence ratings

**DOI:** 10.1101/068601

**Authors:** Stephen M. Fleming

## Abstract

Metacognition refers to the ability to reflect on and monitor one’s cognitive processes, such as perception, memory and decision-making. Metacognition is often assessed in the lab by whether an observer’s confidence ratings are predictive of objective success, but simple correlations between performance and confidence are susceptible to undesirable influences such as response biases. Recently an alternative approach to measuring metacognition has been developed (Maniscalco & Lau, 2012) that characterises metacognitive sensitivity (meta-*d*′) by assuming a generative model of confidence within the framework of signal detection theory. However, current estimation routines require an abundance of confidence rating data to recover robust parameters, and only provide point estimates of meta-*d*’. In contrast, hierarchical Bayesian estimation methods provide opportunities to enhance statistical power, incorporate uncertainty in group-level parameter estimates and avoid edge-correction confounds. Here I introduce such a method for estimating metacognitive efficiency (meta-*d*’/*d*’) from confidence ratings and demonstrate its application for assessing group differences. A tutorial is provided on both the meta-*d*’ model and the preparation of behavioural data for model fitting. Through numerical simulations I show that a hierarchical approach outperforms alternative fitting methods in situations where limited data are available, such as when quantifying metacognition in patient populations. In addition, the model may be flexibly expanded to estimate parameters encoding other influences on metacognitive efficiency. MATLAB software and documentation for implementing hierarchical meta-*d*’ estimation (HMeta-d) can be downloaded at https://github.com/smfleming/HMeta-d.

## INTRODUCTION

Metacognition is defined as “knowledge of one's own cognitive processes” (Flavell, 1979). For example, we can reflect on whether a particular percept is accurate or inaccurate, and this ability to “know that we know” is a central aspect of conscious experience (Schooler, 2002). Consider blindsight, a neurological condition that sometimes arises following selective lesions to primary visual cortex (Weiskrantz et al., 1974). A blindsight patient may perform a task (e.g. discriminating the location of a stimulus) at a reasonably high level in the otherwise blind field, and yet lack insight as to whether they have performed accurately on any given trial (Persaud et al., 2007). It is plausible that a joint lack of metacognition and conscious visual experience are both consequences of disruptions to higher-order representations (Lau & Rosenthal, 2011; Ko & Lau, 2012). While there are clearly other drivers of confidence in one's task performance aside from sensory uncertainty (such as response requirements; Pouget et al., 2016; Denison, in press), understanding the mechanisms supporting metacognition may shed light on the putative underpinnings of conscious experience. Understanding the relationship between metacognition and perceptual and cognitive processes also has broader application in work on judgment and decision-making (Lichtenstein, Fischhoff, & Phillips, 1982), developmental psychology (Goupil, Romand-Monnier, & Kouider, 2016; L. G. Weil et al., 2013), social psychology (Heatherton, 2011) and clinical disorders (David, Bedford, Wiffen, & Gilleen, 2012; Moeller & Goldstein, 2014).

Metacognitive *sensitivity* can be assessed by the extent to which an observer's confidence ratings are predictive of their actual success. Consider a simple decision task such as whether a briefly flashed visual stimulus is categorised as being tilted to the left or right, followed by a confidence rating in being correct. The task of assessing response accuracy using confidence ratings is often called the “type 2 task” (Clarke, Birdsall, & Tanner, 1959; Galvin, Podd, Drga, & Whitmore, 2003) to differentiate it from the “type 1 task” of discriminating between states of the world (e.g. left or right tilts). If higher confidence ratings are given after correct judgments and lower confidence ratings after incorrect judgments, we can ascribe high metacognitive sensitivity to the subject. Thus a simple and intuitive way of assessing metacognitive sensitivity is to correlate confidence with accuracy (Nelson, 1984).

However, confidence-accuracy correlations (e.g. gamma and phi correlations) are affected by the confounding factors of type 1 performance (*d’*) and type 2 response bias (overall level of confidence; Fleming & Lau, 2014; Masson & Rotello, 2009). Consider two subjects A and B performing the same task but with different baseline levels of performance. A and B may have the same underlying metacognitive ability, but their confidence-accuracy correlations may differ due to differing performance levels. In this situation we may erroneously conclude that A and B have different metacognition, despite their underlying metacognitive ability being equal. More generally, an important lesson from the signal detection theory (SDT) approach to modelling type 1 and type 2 tasks is that type 1 sensitivity (*d’*) and type 1 criterion (*c*) influence measures of type 2 sensitivity (Galvin et al., 2003).

Recently an alternative approach to measuring metacognitive sensitivity has been developed by Maniscalco & Lau (2012). This approach posits a generative model of confidence reports within the framework of signal detection theory (Figure 1A). Fitting the model to data returns a parameter, meta-*d’*, that reflects an individual's metacognitive sensitivity. Specifically, meta-*d’* is the value of type 1 performance (*d’*) that would have been predicted to give rise to the observed confidence rating data assuming an ideal observer with type 1 *d’* = meta-*d’*. Meta-*d’* can then be compared with actual *d’* and a relative measure of metacognitive sensitivity can then be calculated as a ratio (*meta-d’/d’*) or subtraction (meta-*d’*-*d’*). Meta-*d’*/*d’* is a measure of *metacognitive efficiency* – given a particular level of task performance, how efficient is the individual’s metacognition? If meta-*d’* = *d’*, then the observer is metacognitively “ideal”, using all the information available for the type 1 task when reporting type 2 confidence. However, we might find that meta-*d’* < *d’*, due to some degree of noise or imprecision introduced when rating one’s confidence. Conversely we may find that meta-*d’* > *d’* if subjects are able to draw on additional information such as hunches (Rausch & Zeheleitner, 2016; Scott, Dienes, Barrett, Bor, & Seth, 2014) further processing of stimulus information (Charles, Van Opstal, Marti, & Dehaene, 2013; Rabbitt & Vyas, 1981) or knowledge of other influences on task performance when making their metacognitive judgments (Fleming & Daw, 2017).

**Figure 1.**
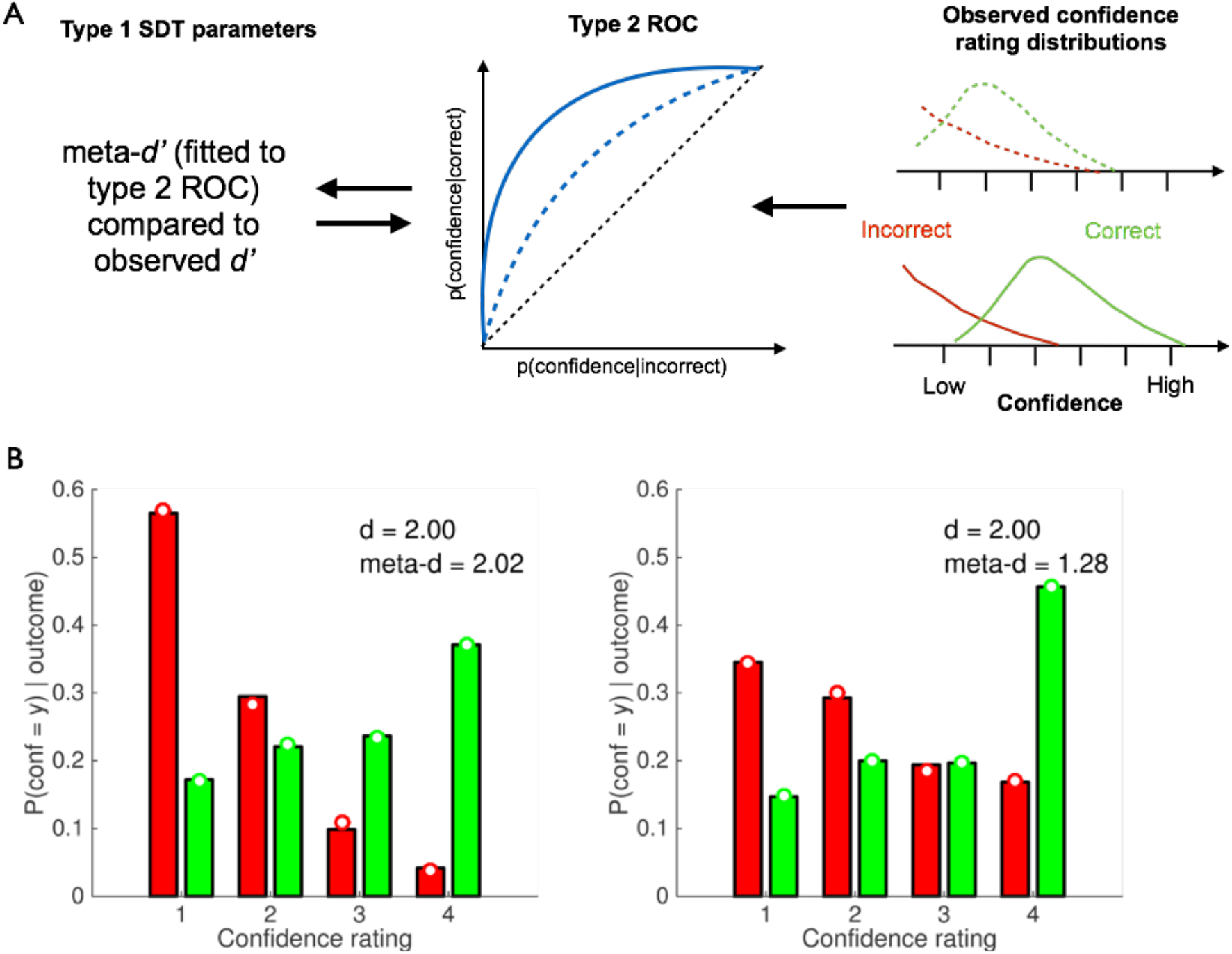
The meta-d’ model. A) The right-hand panel shows schematic confidence-rating distributions conditional on correct and incorrect decisions. A subject with good metacognitive sensitivity will provide higher confidence ratings when they are correct, and lower ratings when incorrect, and these distributions will only weakly overlap (solid lines). Conversely a subject with poorer metacognitive sensitivity will show greater overlap between these distributions (dotted lines). These theoretical correct/error distributions are obtained by “folding” a type 1 SDT model around the criterion (see Galvin et al., 2003, for further details), and normalising such that the area under each curve sums to 1. The overlap between distributions can be calculated through type 2 receiver operating characteristic (ROC) analysis (middle panel). The theoretical type 2 ROC is completely determined by an equal-variance Gaussian signal detection theory model; we can therefore invert the model to determine the type 1 *d’* that best fits the observed confidence rating data, which is labelled meta-*d’*. Meta-*d’* can be directly compared to the type 1 *d’* calculated from the subject’s decisions – if meta-*d’* is equal to *d’*, then the subject approximates the ideal SDT prediction of metacognitive sensitivity. B) Simulated data from a SDT model with *d’* = 2. The y-axis plots the conditional probability of a particular rating given the first-order response is correct (green) or incorrect (red). In the right-hand panel, Gaussian noise has been added to the internal state underpinning the confidence rating (but not the decision) leading to a blurring of the correct/incorrect distributions. Open circles show fits of the meta-*d’* model to each simulated dataset.

The properties of the meta-*d’* model have been thoroughly explored in previous articles (Barrett, Dienes, & Seth, 2013; Fleming & Lau, 2014; Maniscalco & Lau, 2014; 2012). The goal of the present paper is twofold. First, I introduce a new method for estimating meta-*d’*/*d’* from confidence ratings using hierarchical Bayes, and provide a tutorial on its usage. Second, I demonstrate the benefits of applying this method to derive group-level estimates of metacognitive efficiency in situations where data are limited.

Previously *meta-d’* has been fitted using gradient descent on the likelihood (maximum likelihood estimation; MLE), sum-of-squared error (SSE) or using analytic approximation (Barrett et al., 2013; Maniscalco & Lau, 2012). However, several factors make a Bayesian approach attractive for typical metacognition studies:

1. Point estimates of meta-*d’* are inevitably noisy. Several parameters must be estimated in the signal detection model, including multiple type 2 criteria (specifically, (*k* − 1)×2, where *k* = number of confidence ratings available). One common issue in cognitive neuroscience is that trial numbers per condition are also low (e.g. in patient studies, or tasks conducted in conjunction with neuroimaging), and frequentist estimates of hit and false-alarm rates fail to account for uncertainty about these rates that is a consequence of finite data. A Bayesian analysis incorporates such uncertainty into parameter estimates.
2. A hierarchical Bayesian approach is the correct way to combine information about within-and between-subject uncertainty. In a typical study, the metacognitive sensitivities of two groups (e.g. patients and controls) are compared. Single-subject maximum likelihood fits are carried out, and the fitted meta-*d’* parameters are entered into an independent samples t-test. Any information about the uncertainty in each subject’s parameter fits is discarded in this procedure. In contrast, using hierarchical Bayes, information about uncertainty is retained, such that group-level parameters are less influenced by single-subject fits that have a high degree of uncertainty. In turn, hierarchical model fits are able to capitalize on the statistical strength offered by the degree to which subjects are similar with respect to one or more model parameters, mutually constraining the subject-level model fits.
3. In fitting SDT models to data, padding (edge correction) is often applied to avoid zero counts of confidence ratings in particular cells (e.g. high confidence error trials; Hautus, 1995; Macmillan & Creelman, 2005). This padding may bias subject-specific parameter estimates particularly when the overall trial number is low. A Bayesian approach avoids the need for edge correction as the generative multinomial model naturally handles zero cell counts, and a hierarchical specification pools data over subjects (Lee, 2008).
4. A hierarchical model makes testing group-level hypotheses natural and straightforward. For example, say we are interested in testing whether a particular patient group has lower metacognitive sensitivity compared to controls. Hierarchical Bayes allows us to directly estimate the posterior distribution of a parameter that characterises the differences between groups, and provides a principled framework for hypothesis testing. Finally, a Bayesian framework for cognitive modelling enjoys other advantages that have been outlined in detailed elsewhere (Kruschke, 2014; Lee & Wagenmakers, 2014). Briefly, they include the ability to gain evidence in favour of the null hypothesis as well as against it; the ability to combine prior information (for example, a prior on the distribution of metacognitive sensitivity in a healthy population) with new data; and the flexible extension of the model to estimate subject-and trial-level influences on metacognition.

The basics of Bayesian estimation of cognitive models are intuitive. First, prior information is specified in the form of probability distributions over model parameters, and observed data are used to update beliefs to construct a posterior distribution or belief in a particular parameter. The “hierarchical” component of hierarchical Bayes simply indicates that multiple instances of a particular parameter (for example, across different subjects) are estimated in the same model. The development of efficient sampling routines for arbitrary models such as Markov chain Monte Carlo (MCMC), their inclusion in freely available software packages such as JAGS (http://mcmc-jags.sourceforge.net; last accessed 31^st^ August 2016) and STAN (http://mc-stan.org; last accessed 31^st^ August 2016) and advances in computing power means that Bayesian estimation of arbitrary models is now straightforward to implement in practice (Kruschke, 2014).

In this paper I briefly introduce the meta-*d’* model and its hierarchical Bayesian variant (further details of the model can be found in the Appendix and in Maniscalco & Lau, 2014). I then provide a step-by-step MATLAB tutorial for fitting meta-*d’* to single-subject and group data. Finally, I conduct parameter recovery simulations to compare hierarchical Bayesian and standard estimation routines. These results show that, particularly when data are limited, the new HMeta-d method outperforms traditional fitting procedures and provides appropriate control over false positives. Model code and examples are freely available online at https://github.com/smfleming/HMeta-d (last accessed 4th January 2017).

## METHODS

### Outline of the meta-d’ model

The meta-*d’* model is summarized in graphical form in Figure 1A. The raw data for the model fit is the observed distribution of confidence ratings conditional on whether a decision is correct or incorrect. Intuitively, if a subject has greater metacognitive sensitivity, they are able to monitor their decision performance by providing higher confidence ratings when they are correct, and lower ratings when incorrect, and these distributions will only weakly overlap (solid lines). Conversely a subject with poorer metacognitive sensitivity will show greater overlap between these distributions (dotted lines). The overlap between distributions can be calculated through type 2 receiver operating characteristic (ROC) analysis. The conditional probability P(confidence = y | accuracy) is calculated for each confidence level; cumulating these conditional probabilities and plotting them against each other produces the type 2 ROC function. A type 2 ROC that bows sharply upwards indicates a high degree of sensitivity to correct/incorrect decisions; a type 2 ROC closer to the major diagonal indicates weaker metacognitive sensitivity.

The area under the type 2 ROC (AUROC2) is itself a useful non-parametric measure of metacognitive sensitivity, indicating how well an observer’s ratings discriminate between correct and incorrect decisions. However, as outlined in the introduction, AUROC2 is affected by type 1 performance. In other words, a change in task performance (*d’* or criterion) is expected, *a priori*, to lead to changes in AUROC2 despite endogenous metacognitive efficiency remaining unchanged. By explicitly modelling the connection between performance and metacognition we can appropriately handle this confound. The core idea behind the meta-*d’* approach is that a single theoretical type 2 ROC is completely determined by an equal-variance Gaussian signal detection theory model with parameters *d’*, criterion *c* and confidence criteria *c_2_* (the arrow going from left to right in Fig. 1A). The converse is therefore also true: an observed type 2 ROC implies a particular type 1 *d’* (the arrow going from right to left in Fig. 1A), conditional on fixing the type 1 criterion *c*, which in the meta-*d’* model is typically set to the observed value. We can then invert the model to determine the type 1 *d’* that best fits the observed confidence rating data. As this pseudo-*d’* is fit only to confidence rating data, and not the subject’s decisions, we label it meta-*d’*. Meta-*d’* can be directly compared to the type 1 *d’* calculated from the subject’s decisions – if meta-*d’* is equal to *d’*, then the subject approximates the ideal SDT prediction of metacognitive sensitivity. The relative values of *d’* and meta-*d’* thus quantify the relative sensitivity of decisions and confidence ratings respectively. A ratio of these quantities (meta-*d’*/*d’*) provides a summary measure of “metacognitive efficiency”.

Figure 1B provides a concrete example. The data in both panels are simulated from a SDT model with *d’* = 2 and symmetric flanking confidence criteria positioned such that stronger internal signals lead to higher confidence ratings on a 1-4 scale. The y-axis plots the conditional probability of a particular rating given the first-order response is correct (green) or incorrect (red). In both panels, the simulations return higher confidence ratings more often on correct trials and lower confidence more often on incorrect trials. However, in the right-hand panel, Gaussian noise has been added to the internal state underpinning the confidence rating (but not the decision). This leads to a blurring of the correct/incorrect distributions, such that higher confidence ratings are used even when the decision is incorrect. The open circles show fits of the meta-d’ model to each simulated dataset. While both fits return type 1 *d’* values of 2.0, the meta-*d’* value in the right-hand panel is much lower than on the left, leading to a meta-*d’*/*d’* ratio of ~64% of optimal. Notably meta-*d’* in the left panel is similar to *d’*, as expected if confidence ratings are generated from an ideal observer model without any additional noise. This example illustrates how meta-*d’* can appropriately recover changes in the fidelity of confidence ratings independently of changes in performance.

### Single-subject optimisation of meta-d’

I first briefly review the standard meta-*d’* model and the maximum likelihood method for obtaining single-subject parameter estimates. The model contains free parameters for meta-*d’* and the positions of the (*k* − 1)×2 confidence criteria, where *k* = number of confidence ratings available. These criteria are response-conditional, with *k*-1 criteria following an S1 response and *k*-1 criteria following an S2 response (*c_S_*_1_ and *c_S_*_2_). The raw data comprise counts of confidence ratings conditional on both the stimulus category (S1 or S2) and response (S1 or S2). Type 1 criterion *c* and sensitivity *d’* are estimated from the data using standard formulae (Macmillan & Creelman, 2005)^1^.

The fitting of meta-*d’* rests on calculating the likelihood of the confidence rating data given a particular type 2 ROC generated by systematic variation of type 1 SDT parameters *d’* and *c*, and type 2 criteria *c_2_*. By convention, the prefix “meta-“ is added to each type 1 SDT parameter in order to indicate that the parameter is being used to fit type 2 ROC curves. Thus, the type 1 SDT parameters *d’ c*, and *c_2_*, when used to characterize type 2 ROC curves, are named meta-*d’*, meta-*c*, and meta-*c_2_*. Describing the observed type 2 ROC in terms of these type 1 SDT parameters underpins the meta-*d’* model.

The Appendix contains equations for deriving type 2 probabilities from the type 1 SDT model for both S1 and S2 responses. Given a particular setting of the parameters meta-*d’*, meta-*c* and meta-*c_2_* these equations specify a multinomial probability distribution *P*(*conf = y* | *stim = i, resp = j*) over observed confidence counts. The likelihood of the type 2 confidence data for a particular setting of parameters *θ* can be characterized using the multinomial model as:

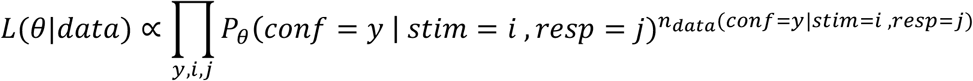

Best-fitting parameters are then obtained by finding parameter settings that maximise the likelihood of the data:

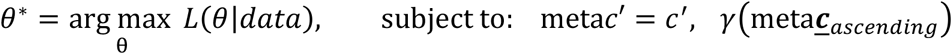

where ***γ*** (*meta****c̱***_*ascending*_)is a Boolean function which returns a value of “true” only if the type 1 and type 2 criteria stand in appropriate ordinal relationships, i.e. each element in ***c̱***_*ascending*_ is at least as large as the previous element, and *c′* is a measure of type 1 response bias.

### Hierarchical Bayesian estimation of meta-d’

In hierarchical Bayesian estimation of meta-*d’* (HMeta-d), the model is similar except group-level prior densities are specified over each of the subject-level parameters referred to in the previous section. A further difference between HMeta-d and single-subject estimation is that the group-level parameter of interest is the ratio meta-*d’*/*d’* rather than meta-*d’* itself. The rationale for this modelling choice is that while each subject or group may differ in type 1 *d’*, our parameter of interest is metacognitive efficiency at the group level, not meta-*d’* (which itself will be influenced by subject-or group-level variability in *d’*). Thus *d’* is treated as a subject-level nuisance parameter^2^. An advantage of this scheme is that group-level inference is carried out directly on metacognitive efficiency rather than a transformed parameter. I specified model parameters such that the prior on log(meta-*d’*/*d’*) encompassed 167 MLE parameter estimates aggregated from behavioural studies of metacognition of perceptual decision-making in our laboratory (Fleming, Huijgen, & Dolan, 2012; Fleming, Weil, Nagy, Dolan, & Rees, 2010; Palmer, David, & Fleming, 2014; L. G. Weil et al., 2013; Figure 2B). This prior was chosen to roughly capture the shape of the empirical distribution, while allowing additional variance in order to relax its influence on posterior estimates. The priors on both log(meta-*d’*/*d’*) and type 2 criteria weakly constrain parameter values to sensible ranges, and can be easily changed by the user in the model specification files. A log-normal prior is appropriate for a ratio parameter, ensuring that increases and decreases relative to the expected value of 1 are given equal weight (Howell, 2009; Keene, 1995).

**Figure 2.**
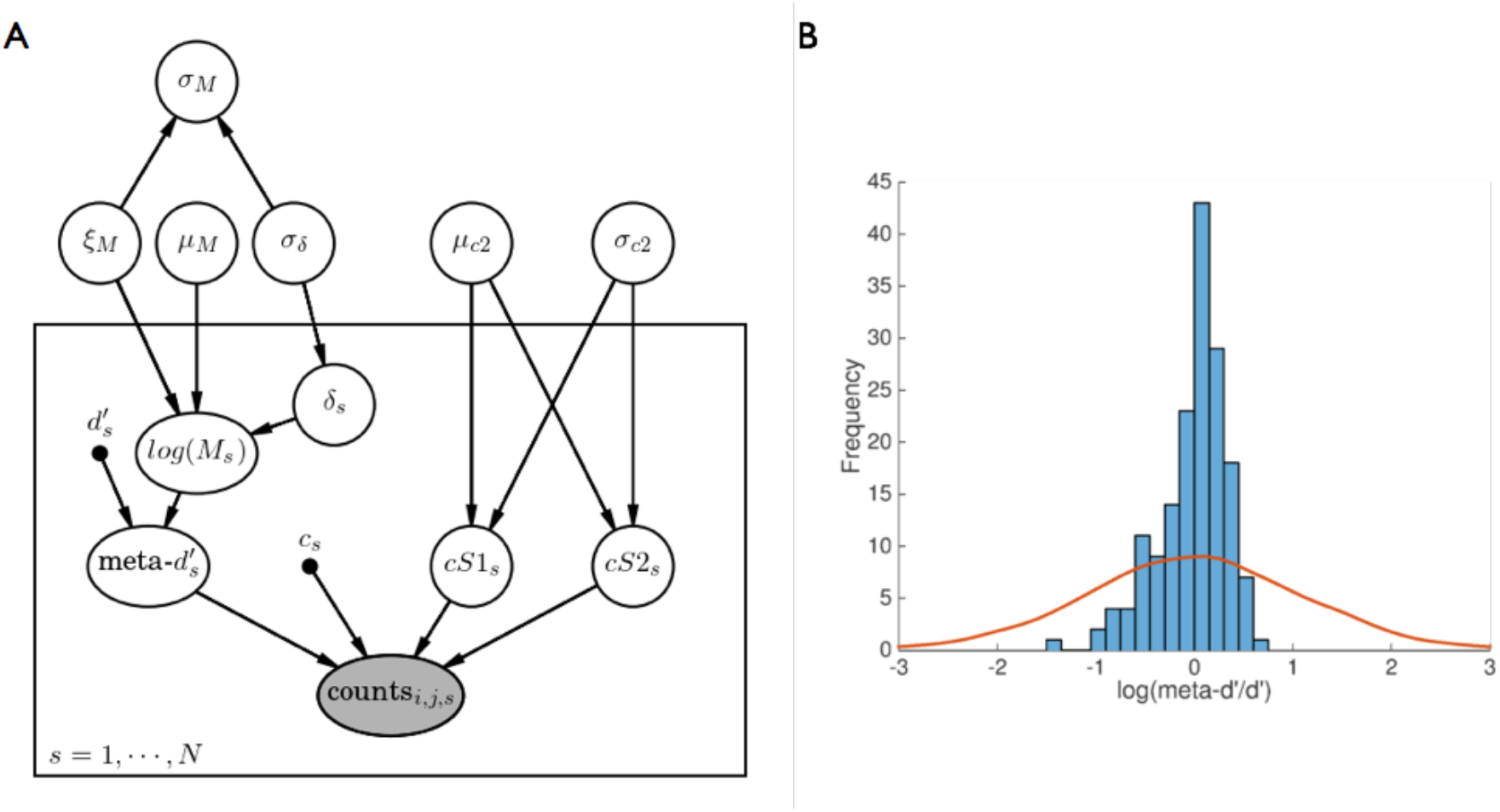
The hierarchical meta-d’ model. A) Probabilistic graphical model for estimating metacognitive efficiency using hierarchical Bayes (*HMeta-d*). The nodes represent all the relevant variables for parameter estimation, and the graph structure is used to indicate dependencies between the variables as indicated by directed arrows. As is convention, unobserved variables are represented without shading and observed variables (in this case, confidence rating counts) are represented with shading. Point estimates for type 1 *d’* and criterion are represented as black dots, and the box encloses participant-level parameters subscripted with s. The main text contains a description of each node and its prior distribution. Figure created using the Daft package in Python (http://daft-pgm.org; last accessed 31^st^ August 2016). B) Prior over the group-level estimate of log(meta-*d’*/*d’*) (μ_M_). The solid line shows a kernel density estimate of samples from the prior; the histogram represents empirical meta-*d’*/*d’* estimates obtained from 167 subjects (see main text for details).

Dependencies between nodes in the HMeta-d model are illustrated as a probabilistic graphical model in Figure 2A. The box encloses participant-level parameters subscripted with *s*. Each node is specified as follows (where *M* denotes log(meta-*d’*/*d’*)):

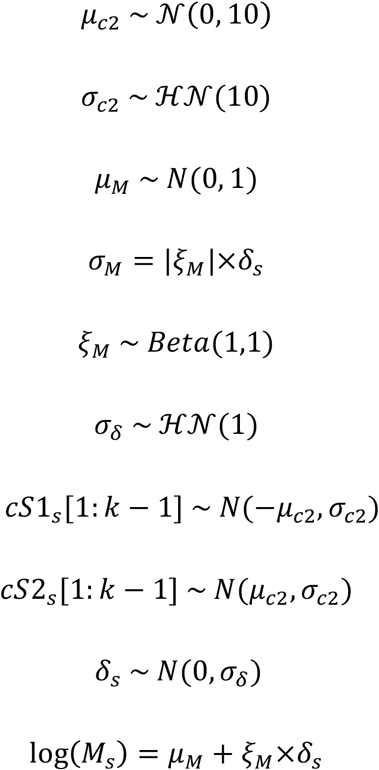

𝒩 represents a normal distribution parmeterised by mean and standard deviation; 𝓗𝒩 represents a positive-only, half-normal parameterised by standard deviation. *μ* and *σ* represent the group-level prior means and standard deviations of subject-level parameters. Thus *μ_c2_* and *σ_c2_* refer to the mean and SD of the type 2 criteria, and *μ_M_* and *σ_M_* to the mean and SD of log(meta-*d’*/*d’*). During model development, it was observed that the hierarchical variance parameter *σ_M_* occasionally became “trapped” near zero during sampling. This problem is fairly common in hierarchical models, and one solution is parameter expansion, whereby the original model is augmented by redundant multiplicative parameters that introduce an additional random component in the sampling process (Gelman & Hill, 2007; Lee & Wagenmakers, 2014). Here I employ the scheme suggested by Matzke, Lee & Wagenmakers (2014), such that the mean and variance of log(*M_s_*) are scaled by a redundant multiplicative parameter *ξ_M_*. The posterior on *σ_M_* can then be recovered by adjusting for the influence of this additional random component.

The HMeta-d toolbox uses Markov chain Monte Carlo (MCMC) sampling as implemented in JAGS (Plummer, 2003) to estimate the joint posterior distribution of all model parameters, given the model specification and the data. This estimation takes the form of samples from the posterior, with the entire sequence of samples known as a chain. It is important to check that these samples approximate the “stationary distribution” of the posterior; i.e. that they are not affected by the starting point of the chain(s), and the sampling behaviour is roughly constant over time without slow drifts or autocorrelation. The default settings of the toolbox discard early samples to avoid sensitivity to initial values and run multiple chains, allowing the user to diagnose convergence problems as described below.

### Preparing confidence rating data

Fitting of group-level data in the HMeta-d toolbox requires similar data preparation to that required when obtaining single-subject fits using MLE or SSE in Maniscalco & Lau’s MATLAB code (available at http://www.columbia.edu/~bsm2105/type2sdt/; last accessed 31^st^ August 2016). I therefore start with a short tutorial on preparing data for estimating single-subject meta-*d’*, before explaining how to input data from a group of subjects into the hierarchical model.

Data from each subject need to be coerced into two vectors, `nR_s1` and `nR_s2`, which contain confidence-rating counts for when the *stimulus* was S1 and S2, respectively. Each vector has length *k* × 2, where *k* is the number of ratings available. Confidence counts are entered such that the first entry refers to counts of maximum confidence in an S1 response, and the last entry to maximum confidence in an S2 response. For example, if three levels of confidence rating were available and `nR_s1 = [100 50 20 10 5 1]`, this corresponds to the following rating counts following S1 presentation:

responded S1, rating=3 : 100 times
responded S1, rating=2 : 50 times
responded S1, rating=1 : 20 times
responded S2, rating=1 : 10 times
responded S2, rating=2 : 5 times
responded S2, rating=3 : 1 time

This pattern of responses corresponds to responding “high confidence, S1” most often following S1 presentations, and least often with “high confidence, S2”. A mirror image of this vector would be expected for `nR_s2`. For example, `nR_s2 = [3 7 8 12 27 89]` corresponds to the following rating counts following S2 presentation:

responded S1, rating=3 : 3 times
responded S1, rating=2 : 7 times
responded S1, rating=1 : 8 times
responded S2, rating=1 : 12 times
responded S2, rating=2 : 27 times
responded S2, rating=3 : 89 times

Together these vectors specify the confidence × stimulus × response matrix that is the basis of the meta-*d’* fit, and can be passed directly into Maniscalco & Lau’s `fit_meta_d_MLE` function to estimate meta-*d’* on a subject-by-subject basis.

### Fitting a hierarchical model

Estimating a group-level model using HMeta-d requires very little extra work. In HMeta-d, the `nR_s1` and `nR_s2` variables are cell arrays of vectors, with each entry in the cell containing confidence counts for a single subject. For example, to specify the confidence counts following S1 presentation listed above for subject 1, one would enter in MATLAB:
`nR_S1{1} = [100 50 20 10 5 1]`
and so on for each subject in the dataset. These cell arrays then contain confidence counts for all subjects, and are passed in one step to the main HMeta-d function:
`fit = fit_meta_d_mcmc_group(nR_S1, nR_S2)`

An optional third argument to this function is `mcmc_params` which is a structure containing flags for choosing different model variants, and for specifying the details of the MCMC routine. If omitted reasonable default settings are chosen.

The call to `fit_meta_d_mcmc_group` returns a “fit” structure with several subfields. The key parameter of interest is `fit.mu_logMratio`, which is the mean of the posterior distribution of the group-level log(meta-*d’*/*d’*). `fit.mcmc` contains the samples of each parameter, which can be plotted with the helper function `plotSamples`. For instance to plot the MCMC samples of *μ_M_*, one would enter:
`plotSamples(exp(fit.mcmc.samples.mu_logMratio))`

Note the “exp” to allow plotting of meta-*d’*/*d’* rather than log(meta-*d’*/*d’*). The `exampleFit_` scripts in the toolbox provide other examples, such as how to set up response-conditional models and to visualise subject-level fits.

An important step in model fitting is checking that the MCMC chains have converged to a stationary distribution. While there is no way to guarantee convergence for a given number of MCMC samples, some heuristics can help identify problems. By using `plotsamples`, we can visualise the traces to check that there are no drifts or jumps and that each chain occupies a similar position in parameter space. Another useful statistic is Gelman & Rubin’s scale-reduction statistic *R̂*, which is stored in the field `fit.mcmc.Rhat` for each parameter (Gelman & Rubin, 1992). This provides a formal test of convergence that compares within-chain and between-chain variance of different runs of the same model, and will be close to 1 if the samples of the different chains are similar. Large values of *R̂* indicate convergence problems and values < 1.1 suggest convergence.

As well as obtaining an estimate for group-level meta-*d’/d’*, we are often interested in our certainty in this parameter value. This can be estimated by computing the symmetric 95% credible interval (CI), which is the interval bounded by the 2.5% and 97.5% percentiles of MCMC samples. An alternative formulation is the 95% highest-density interval (HDI), which is the shortest possible interval containing 95% of the MCMC samples, and is not necessarily symmetric (Kruschke, 2014). The helper functions `calc_CI` and `calc_HDI` take as input a vector of samples and return the 95% CI/HDI:
`calc_CI(exp(fit.mcmc.samples.mu_logMratio(:))`

The colon in the brackets selects all samples in the array regardless of their chain of origin. As HMeta-d uses Bayesian estimation it is straightforward to use the group-level posterior density for hypothesis testing. For instance, if the question is whether one group of subjects has greater metacognitive efficiency than a second group, we can ask whether the CI/HDI of the difference overlaps with zero (see “Empirical examples” below for an example of this). However, note that it is incorrect to use the subject-level parameters estimated as part of the hierarchical model in a frequentist test (e.g. a *t*-test); this violates the independence assumption.

In addition to enabling inference on individual parameter distributions, there may be circumstances in which we wish to compare models of different complexity (see Discussion). To enable this, JAGS returns the deviance information criteria (DIC) for each model which is a summary measure of goodness of fit, while penalising for model complexity (Spiegelhalter, Best, Carlin, & Van Der Linde, 2002; lower is better). While DIC is known to be somewhat biased towards models with greater complexity, it is a common method for assessing model fit in hierarchical models. In HMeta-d the DIC for each model can be obtained in `fit.mcmc.dic`.

### Simulations

To assess properties of the model fit and compare alternative fitting procedures, simulated confidence rating data were generated for pre-specified levels of metacognitive efficiency. Type 2 probabilities *P*(*conf = y|stim, resp*) were computed from the equations in the Appendix for particular settings of meta-*d’*, *c* and *c_2_*. These probabilities were then used to generate multinomial response counts using the function mnrnd in MATLAB, where the sample size of each type 1 response class (hits, false alarms, correct rejections and misses) was obtained from a standard type 1 SDT model with criterion *c* and *d’*. This allowed for independent control over *d’* (i.e. the number of hits and false alarms) and meta-*d’* (the response-conditional distribution of confidence ratings). After determining the value of *d’* for each simulation, the relevant value of meta-*d’* could then be chosen to ensure a particular target meta-*d’*/*d’* level. This procedure is implemented in the MATLAB function `metad_sim` included as part of the toolbox.

## RESULTS

### Example fit

Figure 3A shows the output of a typical call to HMeta-d and the resultant posterior samples of the population-level estimate of metacognitive efficiency, *μ_meta−d’/d’_*, ploted with `plotsamples`. The data were generated as 20 simulated subjects, each with 400 trials and four possible confidence levels (confidence criteria *c_2_* = ±[0.5 11.5]; type 1 criterion *c* = 0). For each subject, type 1 *d’* was sampled from a normal distribution *N*(2,0.2), and meta-*d’*/*d’* was fixed at 0.8. The chains show excellent mixing with a modest number of samples (10,000 per chain; *R̂* = 1.000) and the posterior is centred around the ground truth simulated value.

**Figure 3.**
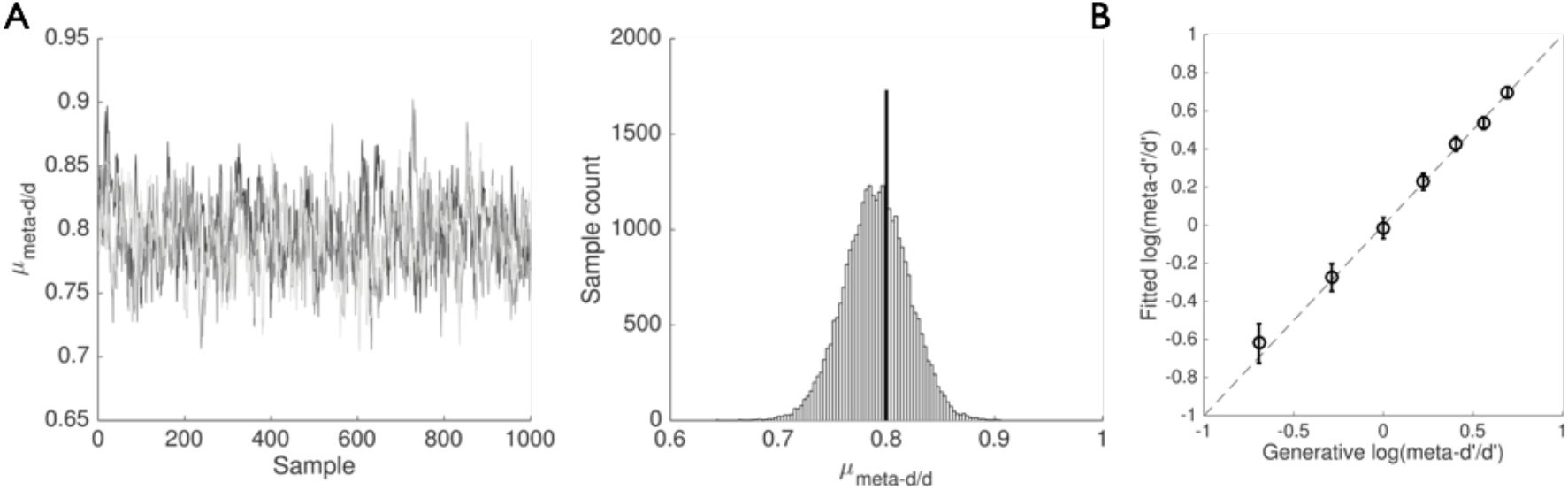
HMeta-d output. A) Example output from HMeta-d fit to simulated data with ground truth *meta*-d’/d’ fixed at 0.8 for 20 subjects. The left panel shows the first 1,000 samples from each of three MCMC chains for parameter *μ_meta-d’/d’_*; the right panel shows all samples aggregated in a histogram. B) Parameter recovery exercise using HMeta-d to fit data simulated from 7 groups of 20 subjects with different levels of meta-*d’*/*d’* = [0.5 0.75 1.0 1.25 1.5 1.75 2]. Error bars denote 95% CI.

### Parameter recovery

To further validate the model a parameter recovery exercise was carried out in which data were simulated from 7 groups of 20 subjects with different levels of meta-*d’*/*d’* = [0.5 0.75 1.0 1.25 1.5 1.75 2]. All other settings were as described in the previous section. Figure 3B plots the fitted group-level *μ_meta−d’/d’_* and its associated 95% CI for each of the simulated datasets against the empirical ground truth, demonstrating robust parameter recovery.

### Empirical examples

To illustrate the practical application of HMeta-d I fit data from a recent experiment that examined metacognitive sensitivity in perceptual and mnemonic tasks in patients with post-surgical lesions and controls (Fleming, Ryu, Golfinos, & Blackmon, 2014). This study found (using single-subject estimates of meta-*d’*/*d’*) that metacognitive efficiency in patients with lesions to anterior prefrontal cortex (aPFC) was selectively compromised on a visual perceptual task but unaffected on a memory task, suggesting that the neural architecture supporting metacognition may comprise domain-specific components differentially affected by neurological insult.

For didactic purposes here I restrict comparison of metacognition in the aPFC patients (N=7) and healthy controls (HC; N=19) on the perceptual task. The task required a two-choice discrimination as to which of two briefly presented patches contained a greater number of small white dots, followed by a continuous confidence rating on a sliding scale from 1 (low confidence) to 6 (high confidence). For analysis these confidence ratings were binned into four quantiles. For each subject confidence rating data (levels 1-4) were sorted according to the position of the target stimulus (L/R) and the subject’s response (L/R), thereby specifying the two nR_S1 and nR_S2 arrays required for estimating meta-*d’*.

For each group I constructed cell arrays of confidence counts and estimated *μ_meta−d′/d’_* with the default settings in HMeta-d. The resultant posterior distributions are plotted in Figure 4A, and the posterior distribution of the difference is shown in Figure 4B. Several features are evident from these outputs. First, there is a reduced metacognitive efficiency in the aPFC group compared to controls, as revealed by the 95% CI of the difference being greater than zero (right-hand panel). Second, the posterior distribution of metacognitive efficiency in the healthy controls is overlapping with the optimal estimate of 1. Finally, for the aPFC group, which compromises fewer subjects, there is a higher degree of uncertainty about the true metacognitive efficiency – the width of the posterior distribution is greater. This is due to the parameter estimate being constrained by fewer data points and is a natural consequence of the Bayesian approach.

**Figure 4.**
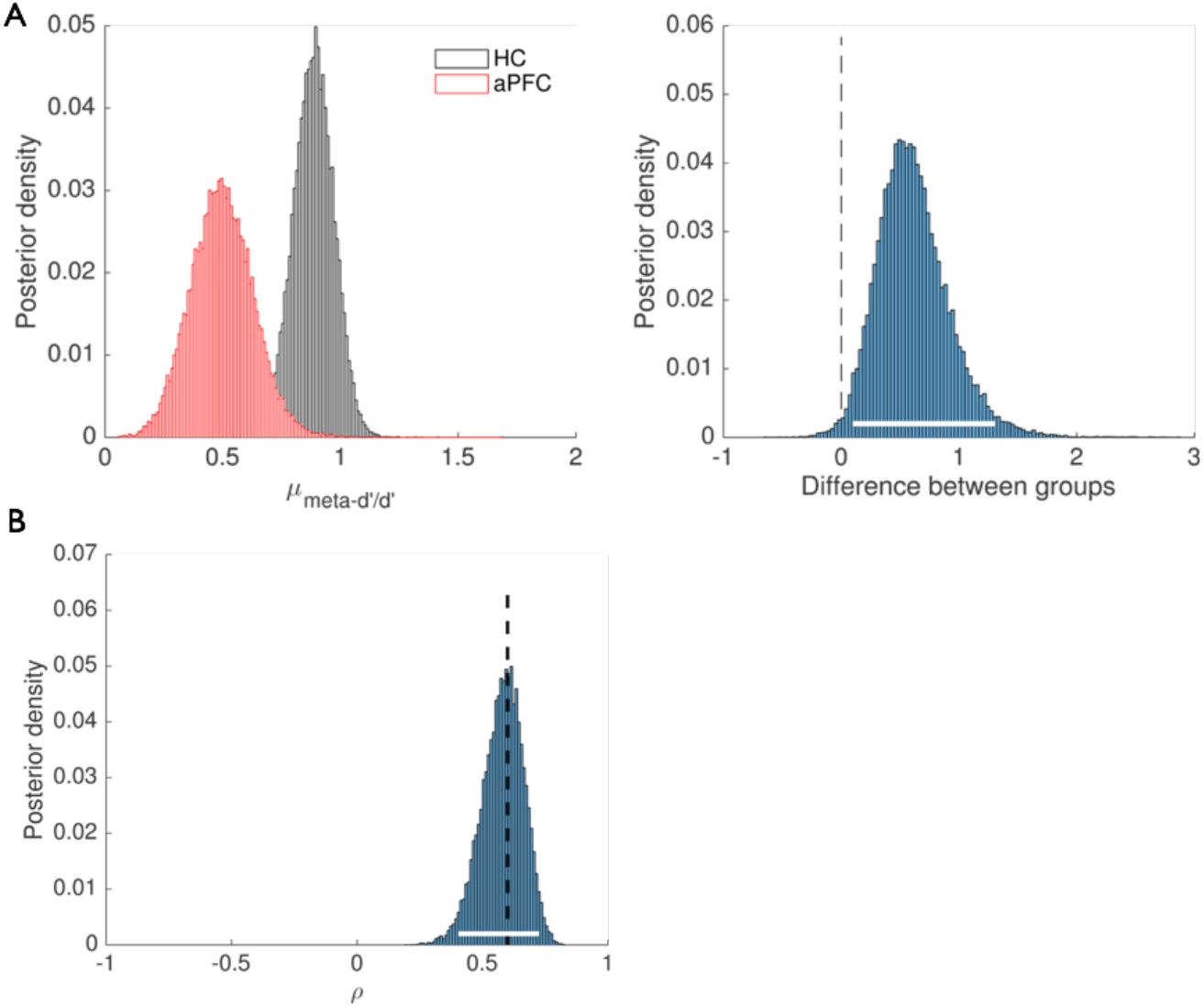
Empirical applications of HMeta-d. A) HMeta-d fits to data from the perceptual metacognition task reported in Fleming et al. (2014). Each histogram represents posterior densities of *μ_meta – d’/d’_* for two groups of subjects: HC = healthy controls; aPFC = anterior prefrontal cortex lesion patients. The right panel shows the difference (in log units) between the group posteriors. The white bar indicates the 95% CI which excludes zero. B) Example of extending the HMeta-d model to estimate the correlation coefficient ρ between metacognitive efficiencies in two domains. The dotted line shows the ground-truth correlation between pairs of meta-d’/d’ values for 100 simulated subjects.

### Comparison of fitting procedures

To compare the quality of the fit of the hierarchical Bayesian method against MLE and SSE point-estimate approaches, I ran a series of simulation experiments to investigate parameter recovery of known meta-*d’*/*d’* ratios for different *d’* and type 2 criteria placements across a range of trial counts.

In each experiment I simulated confidence rating data for groups of N=20 subjects while manipulating the number of trials (20, 50, 100, 200, 400). In the first set of experiments type 1 *d’* was selected from the set (0.5, 1, 2), and two type 2 criteria were specified such that 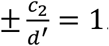. The generated data thus consisted of a 2 (stimulus) × 2 (responses) × 2 (high/low confidence) matrix of response counts. In the second set of experiments type 1 d’ was kept constant at 1, and the type 2 criteria were selected from the set 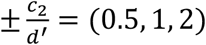. Generative meta-*d’*/*d’* was fixed at 1, and type 1 criterion was fixed at 0.

Each simulated subject’s data was fit using the MLE and SSE routines available from http://www.columbia.edu/~bsm2105/type2sdt/, correcting for zero response counts by adding 0.25 to all cells (a generalisation of the log-linear correction typically applied when estimating type 1 *d’*, as recommended by Hautus (1995)). For each group of 20 subjects the mean meta-*d’*/*d’* ratio and the output of a one-sample *t*-test against the null value of 1 was stored. The same data (without padding) were entered into the hierarchical Bayesian estimation routine as described above and the posterior mean stored. A false positive was recorded if a one-sample *t*-test against the null value (meta-*d’*/*d’* = 1) was significant (*P* < 0.05) for the MLE/SSE approaches, or if the symmetric 95% credible interval excluded 1 for the hierarchical Bayesian approach. This procedure was repeated 100 times for each setting of trial counts and parameters.

Figure 5A and B shows the results of Experiments 1 and 2, respectively, for medium levels of metacognitive efficiency (meta-*d’*/*d’* = 1). For intermediate values of *d’* and criteria (middle panels), all methods perform similarly, and recover the true meta-*d’*/*d’* ratio. However when *d’* is low, or criteria are extreme, the MLE and SSE methods tend to misestimate metacognitive efficiency when the number of trials per subject is < 100, leading to high false positive rates. These misestimations are similar to the effect of zero cell-count corrections on recovery of type 1 *d’* (Hautus, 1995). In contrast, HMeta-d provides accurate parameter recovery in the majority of cases.

**Figure 5.**
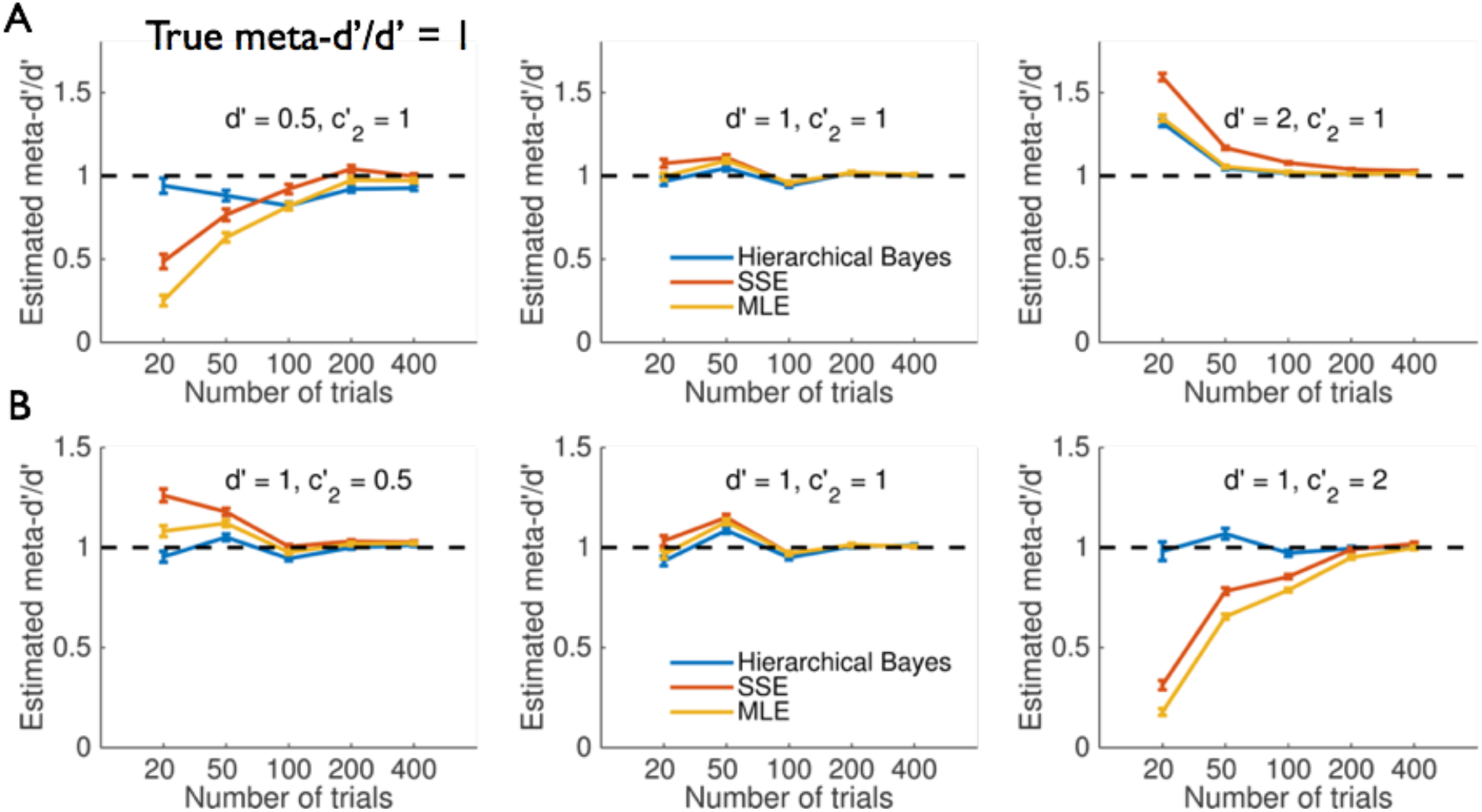
Simulation experiments – medium metacognitive efficiency (meta-d’/d’ = 1). A, B) Estimated meta-d’/d’ ratio for different fitting procedures while varying A) d’ values or B) type 2 criteria placements. Each data point reflects the average of 100 simulations each with N=20 subjects. Error bars reflect standard errors of the mean. The ground truth value of meta-*d’*/*d’* is shown by the dotted line.

Why does HMeta-d outperform classical estimation procedures in this case? There are two possible explanations. First, HMeta-d may be more efficient at retrieving true parameter values, even when trial counts are low, by capitalizing on the hierarchical structure of the model in order to mutually constrain subject-level fits. Alternatively, HMeta-d may rely more on the prior when data are scarce, thus shrinking group estimates to the prior mean. The second explanation predicts that HMeta-d would become less accurate when true metacognitive efficiency deviates from the prior mean (meta-*d’*/*d’* ~ 1).

To adjudicate between these explanations I repeated the simulations at low (meta-*d’*/*d’* = 0.5) and high (meta-*d’*/*d’* = 1.5) metacognitive efficiency (Figures 6 and 7). These results show that HMeta-d is able to retrieve the true meta-*d’*/*d’* even when metacognitive efficiency is appreciably less than or greater than 1 (see also Figure 3B), consistent with the prior exerting limited influence on the results. One notable exception is found when type 1 *d’* is high, and trial counts are very low (~20 per subject); in this case (upper right-hand panels), all fitting methods tend to overestimate metacognitive efficiency.

**Figure 6.**
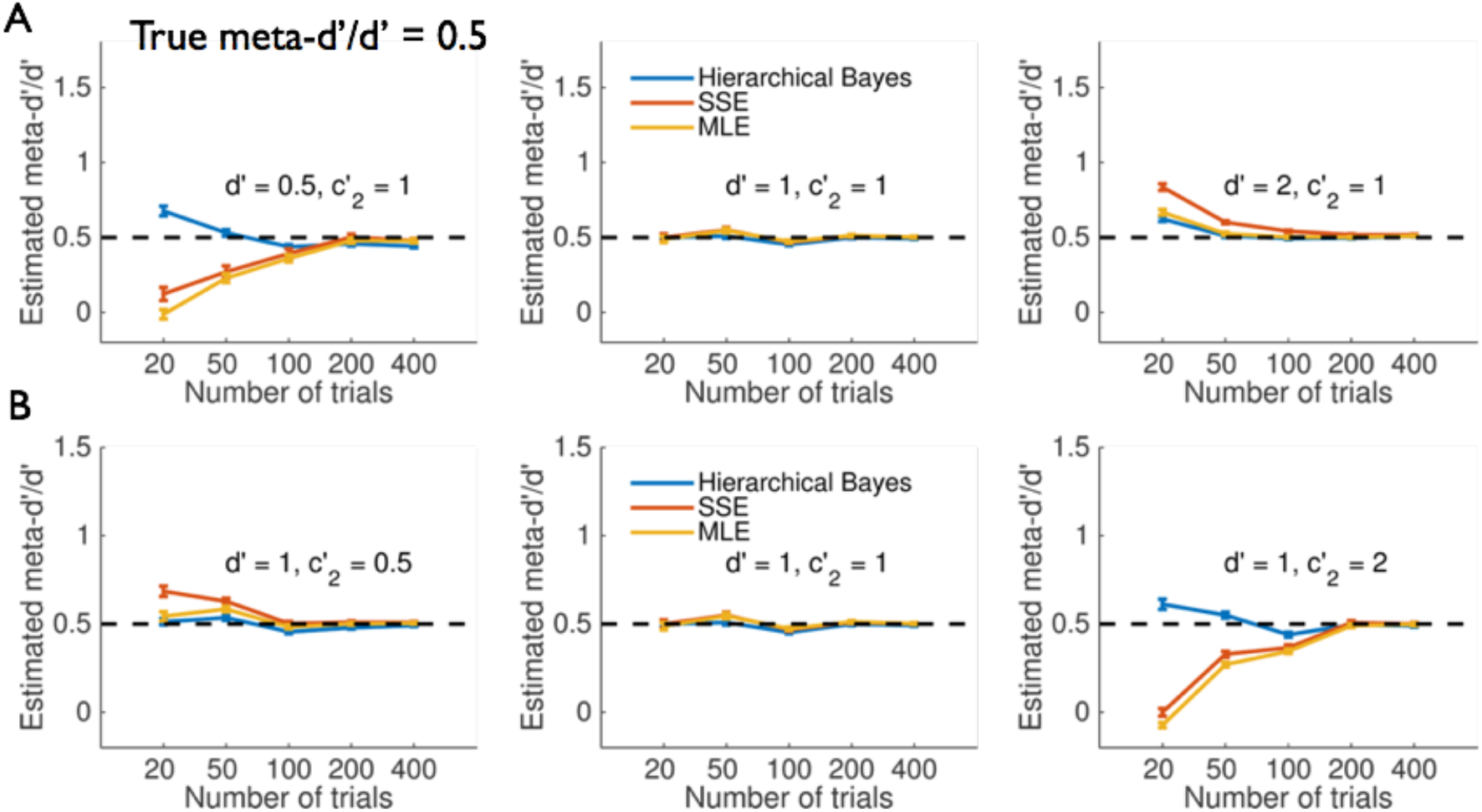
Simulation experiments – low metacognitive efficiency (meta-d’/d’ = 0.5). For legend see Figure 5.

**Figure 7.**
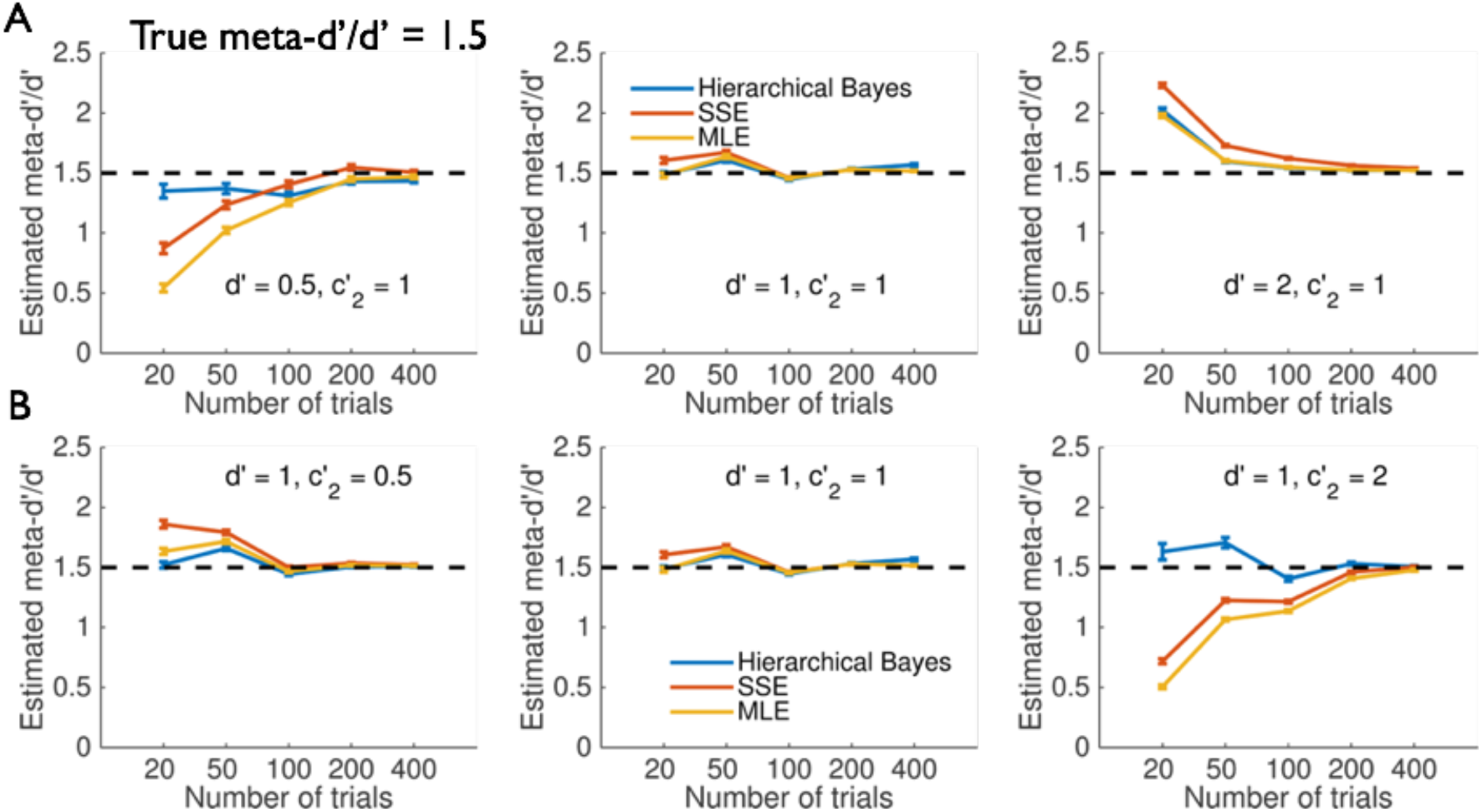
Simulation experiments – high metacognitive efficiency (meta-d’/d’ = 1.5). For legend see Figure 5.

Figure 8 provides a summary of false positive rates recorded across all experiments for the three methods. Point-estimate approaches (SSE and MLE) return unacceptably high false positive rates when trial counts are less than ~200 per subject, due to consistent over-or underestimation of metacognitive efficiency. In contrast, HMeta-d provides good control of the false positive rate in all cases except when trial counts are very low (< 50 per subject).

**Figure 8.**
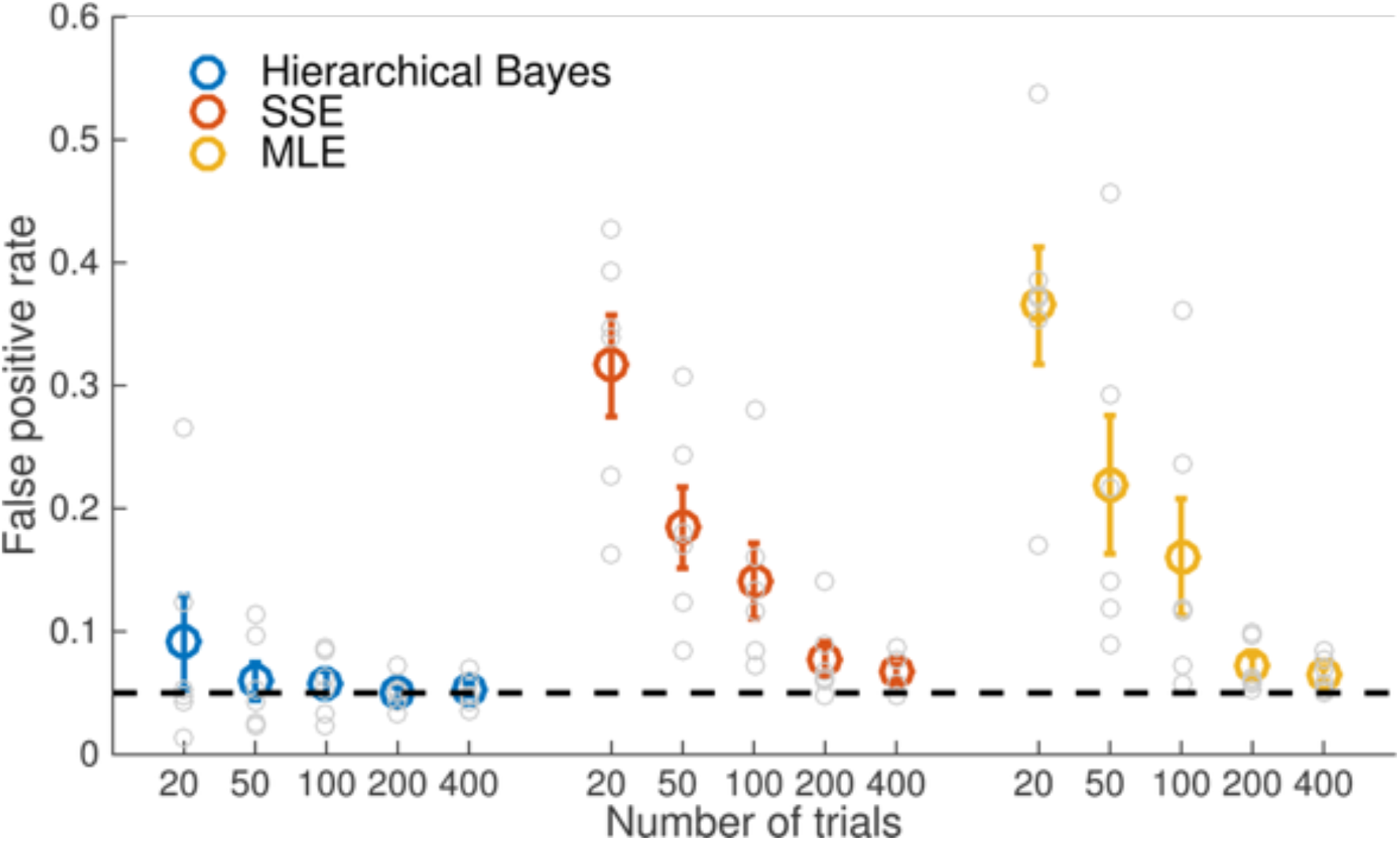
Observed false positive rates for each fitting procedure. Average false positive rates for hypothesis tests against ground truth meta-d’/d’ values from the simulations in Figures 5, 6 and 7. Individual data points reflect single experiments (the false positive rate for a particular combination of metacognitive efficiency level, parameters and trial count). Error bars reflect standard errors of the mean. For trial counts < 200, MLE or SSE methods result in unacceptably high false positive rates due to consistent over-or underestimation of metacognitive efficiency.

### Flexible extensions of the basic model

An advantage of working with Bayesian graphical models is that they are easily extendable to estimate other influences on metacognitive efficiency in the context of the same model (Lee & Wagenmakers, 2014). For instance, one question of interest is whether metacognitive ability in one domain, such as perception, is predictive of metacognitive ability in another domain, such as memory. Evidence pertaining to this question is mixed: some studies have found evidence for a modest correlation in metacognitive efficiency across domains (Ais, Zylberberg, Barttfeld, & Sigman, 2016; McCurdy et al., 2013) whereas others have reported a lack of correlation (Baird, Smallwood, Gorgolewski, & Margulies, 2013; Kelemen, Frost, & Weaver, 2000). One critical issue in testing this hypothesis is that uncertainty in the model’s estimate of meta-*d’* should be incorporated into an assessment of any correlation between the two domains. This is naturally accommodated by embedding an estimate of the correlation coefficient in a hierarchical estimation of metacognitive efficiency.

To expand the model, each subject’s metacognitive efficiencies in the two domains (*M1, M2*) are specified as draws from a bivariate Gaussian^3^:

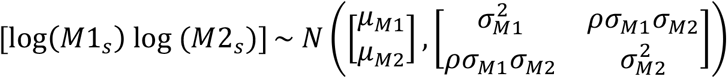

Priors were specified as follows:

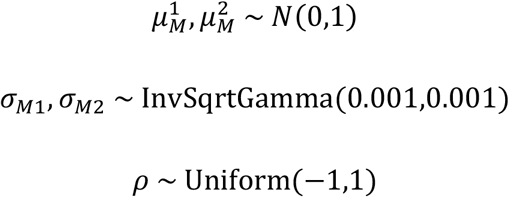

To demonstrate the application of this expanded model I simulated 100 subjects’ confidence data from the type 2 SDT model in two “tasks”. Each task’s generative meta-*d’*/*d’* was drawn from a bivariate Gaussian with mean = *μ_M1_* = *μ_M2_* = 0.8 and standard deviations *σ_M_*_1_ = *σ_M_*_2_ = 0.5. Type 1 *d’* was generated separately for each task from a *N*(2,0.2) distribution. The generative correlation coefficient *ρ* was set to 0.6. Data from both domains are then passed into the model simultaneously, and a group-level posterior distribution on the correlation coefficient *ρ* is returned. Figure 4B shows this posterior together with the 95% CI, which encompasses the generative correlation coefficient.

## DISCUSSION

The quantification of metacognition from confidence ratings is a question with application in several subfields of psychology and neuroscience, including consciousness, decision-making, memory, education, aging, and psychiatric disorders. There are now several tools in the psychologist’s armoury for estimating how closely subjective reports track task performance (Fleming & Lau, 2014). An important advance is the recognition that simple correlation coefficients are affected by fluctuations in performance and confidence bias, and the meta-*d’* model was developed to allow correction of metacognitive sensitivity for these potential confounds (Maniscalco & Lau, 2012).

The hierarchical Bayesian approach to estimating metacognitive efficiency introduced here enjoys several advantages. It naturally incorporates variable uncertainty about finite hit and false-alarm rates; it is the correct way to incorporate information about within-and between-subject uncertainty; it avoids the need for edge correction or data modification, and provides a flexible framework for hypothesis testing and model expansion. The toolbox provides a simple MATLAB implementation that harnesses the MCMC sampler JAGS to return posterior distributions over group-level model parameters. The tutorial outlined how data preparation is identical to that required for the existing maximum-likelihood routines, allowing the user to easily apply both approaches once data are in the correct format. In simulation experiments, the hierarchical approach recovered more accurate parameter estimates than commonly used alternatives (MLE and SSE), and this benefit was greatest when there are limited numbers of trials per subject (Figure 5). It is notable that the point-estimate approaches severely underestimate average meta-d’/d’ ratios for low *d’* and trial numbers < ~100 per subject, leading to a high false positive rate. Given that low (type 1) *d’* is commonplace in psychophysical studies of conscious awareness and metacognition, such biases may lead to erroneous conclusions that metacognitive efficiency is below the ideal observer prediction. In contrast, over-estimations were observed when *d’* was high.

### Practical recommendations for quantifying metacognition

If group-level estimates of meta-*d’/d’* are of primary interest, HMeta-d allows direct, unbiased inference at this upper level of the hierarchy while appropriately handling participant-level uncertainty. The HMeta-d toolbox also allows Bayesian estimation of single-subject meta-*d’*, but if single-subject estimates are of primary interest, the MLE approach may be simpler and computationally less expensive. However advantages of using a Bayesian approach even for single-subject estimates is that uncertainty in parameter estimates can be easily quantified (as the posterior credible interval), with such uncertainty appropriately reducing as trial count increases, and edge correction confounds are avoided.

More generally, whether one should use metacognitive sensitivity (e.g. meta-*d’* or AUROC2) or metacognitive efficiency (meta-*d’*/*d’*) as a measure of metacognition depends on the goal of an analysis. For example, if we are interested in establishing the presence or absence of metacognition in a particular condition, such as when performance is particularly low (Scott et al., 2014) or in particular subject groups such as human infants (Goupil et al., 2016), computing metacognitive sensitivity alone may be sufficient. However, when comparing experimental conditions or groups which may differ systematically in performance, estimating metacognitive efficiency appropriately controls for confounds introduced by type 1 performance and response biases. Note however there are also limitations in the applicability of the meta-*d’* model. First and foremost, the task should be amenable to analysis in a 2-choice SDT framework, as fitting meta-*d’* requires specification of a 2 (stimulus) × 2 (response) × N (confidence rating) matrix. If a task does not conform to these specifications (such as one with N alternative responses) then employing an alternative bias-free measure of metacognitive sensitivity such as the area under the type 2 ROC (AUROC2) may be preferable (Fleming & Lau, 2014). In addition, like all analysis approaches, meta-*d’* assumes a particular generative model of the confidence data that is at best incomplete, and untenable in certain circumstances. For instance, equal variance is specified for S1 and S2 distributions^4^ and stable confidence criteria are assumed which may be at odds with findings of serial adjustments in criteria (Norton et al., 2017; Rahnev, Koizumi, McCurdy, D’Esposito, & Lau, 2015; Treisman, 1984).

More broadly, meta-*d’* is primarily a tool for estimating metacognitive sensitivity, and additional considerations are needed when developing a complete model of confidence (Pouget, Drugowitsch, & Kepecs, 2016; Fleming & Daw, 2017). Recent modelling work has sought to explicitly characterise type 1 and type 2 processes (Jang et al., 2012; Fleming & Daw, 2017; Maniscalco & Lau, 2016), permitting flexible modelling of relationships between performance and metacognition. For instance, in Fleming and Daw’s “second-order” model, an underlying generative model of action is specified, and confidence is formulated as an inference on the model’s probability of being correct, conditioned on both internal states and self-action. These frameworks allow for multiple drivers of metacognitive sensitivity, in contrast to the meta-*d’* model which describes sensitivity only relative to type 1 performance. It is thus useful to view meta-*d’* as complementary to these modelling efforts. Just as *d’* provides a bias-free measure of perceptual sensitivity that may be explained by a number of contributing factors, meta-*d’* provides a bias-free metric for metacognitive sensitivity without commitment to a particular processing architecture.

### Future directions

The HMeta-d model code can be flexibly extended to allow estimation of other influences on metacognitive sensitivity. Here one simple example is explored, the specification of a population-level correlation coefficient relating metacognitive efficiencies across domains. More broadly, it may be possible to specify flexible general linear models linking trial-or subject-level variables to meta-*d’* (Kruschke, 2014). Currently this requires bespoke model specification, but in future work we hope to provide a flexible user interface for the specification of arbitrary models (cf. Wiecki, Sofer, & Frank, 2013). Estimation of single-trial influences on metacognitive efficiency, such as attentional state or brain activity, is a particularly intriguing proposition.

Currently, estimation of meta-*d’* requires many trials, restricting studies of the neural basis of metacognitive efficiency to between-condition or between-subject analyses. Extending the HMeta-d framework to estimate trial-level effects on meta-*d’* may therefore accelerate our understanding of the neural basis of metacognitive efficiency.

Also naturally accommodated in a hierarchical framework is the comparison of different model structures for metacognition within and across tasks. A currently open question is whether metacognition relies on common or distinct processes across different domains, such as perception or memory (Ais et al., 2016; Baird et al., 2013; Fleming et al., 2014; McCurdy et al., 2013). One approach to addressing this question is to specify variants of the HMeta-d model in which different parameters are shared across domains, such as meta-*d’* and/or the confidence criteria. Through model comparison, one could then obtain the model that best accounted for the relationship between metacognitive performance across different domains, and shed light on the common and distinct components.

### Conclusions

This paper introduces a hierarchical Bayesian approach to estimating metacognitive efficiency. This approach has several methodological advantages in comparison to current methods that focus on single-subject point estimates, and may prove particularly beneficial for studies of metacognition in patient populations and cognitive neuroscience experiments where often only limited data are available. More broadly, this framework can be flexibly extended to specify and compare different models of meta-*d’* within a common scheme, thereby advancing our understanding of the neural and computational basis of self-evaluation.

## Appendix

### Type 2 SDT model equations

For a discrete confidence scale ranging from 1 to *k, k* − 1 type 2 criteria are required to rate confidence for each response type. We define type 2 confidence criteria for S1 and S2 responses as:

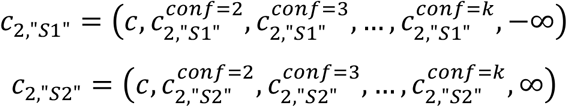

And

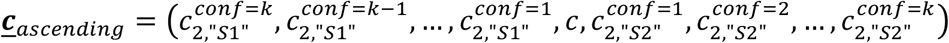

Then the probabilities of each confidence rating conditional on a given stimulus and response are:

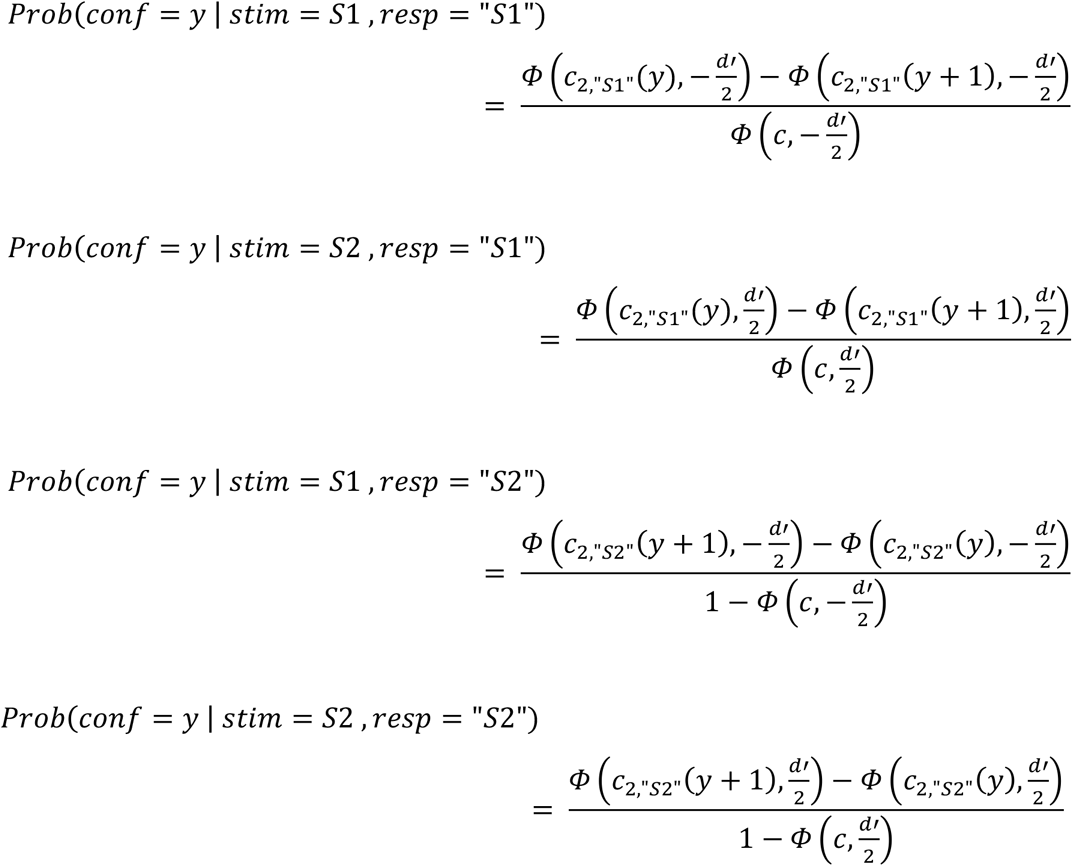

where *Φ*() is the cumulative distribution function of the standard normal distribution.

## Acknowledgements

Funded by a Sir Henry Wellcome Fellowship from the Wellcome Trust (096185) awarded to SMF. The Wellcome Trust Centre for Neuroimaging is supported by core funding from the Wellcome Trust (091593/Z/10/Z).

## Footnotes

1 In HMeta-d there is also a user option for jointly estimating both *d’* and meta-*d’* in a hierarchical framework.

2 Alternative estimation schemes are possible; for instance, calculating the ratio of hierarchical nodes independently encoding meta-*d’* and *d’*. I chose the “nuisance parameter” scheme as it stays closest to the standard MLE approach while directly estimating a group-level node for meta-*d’*/*d’*.

3 Note parameter expansion is omitted here for clarity

4 Maniscalco & Lau’s `fit_meta_d_MLE` code allows setting the ratio of S1 and S2 variances as a free parameter; it would be possible to incorporate a similar parameter in future versions of HMeta-d. However as described by Maniscalco & Lau (2014) there is ambiguity between changes in response specific metacognitive efficiency and the variance ratio, and therefore we recommend users employ the equal-variance model unless they have access to independent estimates of the variance inequality.

## REFERENCES

Ais, J., Zylberberg, A., Barttfeld, P., & Sigman, M. (2016). Individual consistency in the accuracy and distribution of confidence judgments. Cognition, 146, 377–386. http://doi.org/10.1016Zj.cognition.2015.10.006

Baird, B., Smallwood, J., Gorgolewski, K. J., & Margulies, D. S. (2013). Medial and Lateral Networks in Anterior Prefrontal Cortex Support Metacognitive Ability for Memory and Perception. Journal of Neuroscience, 33(42), 16657–16665. http://doi.org/10.1523/JNEUROSCI.0786-13.2013

Barrett, A. B., Dienes, Z., & Seth, A. K. (2013). Measures of metacognition on signal-detection theoretic models. Psychological Methods, 18(4), 535–552. http://doi.org/10.1037/a0033268

Charles, L., Van Opstal, F., Marti, S., & Dehaene, S. (2013). Distinct brain mechanisms for conscious versus subliminal error detection. NeuroImage, 73, 80–94. http://doi.org/10.1016/j.neuroimage.2013.01.054

Clarke, F., Birdsall, T., & Tanner, W. (1959). Two types of ROC curves and definition of parameters. Journal of the Acoustical Society of America, 31, 629–630.

David, A. S., Bedford, N., Wiffen, B., & Gilleen, J. (2012). Failures of metacognition and lack of insight in neuropsychiatric disorders., 367(1594), 1379–1390. http://doi.org/10.1098/rstb.2012.0002

Denison, R.N. (in press) Precision, not confidence, describes the uncertainty of perceptual experience. (Response to John Morrison's “Perceptual confidence”.) Analytic Philosophy.

Flavell, J. H. (1979). Metacognition and cognitive monitoring: A new area of cognitive– developmental inquiry. American Psychologist, 34(10), 906–911. http://doi.org/10.1037/0003-066X.34.10.906

Fleming, S. M., & Daw, N. D. (2017). Self-evaluation of decision-making: A general Bayesian framework for metacognitive computation. Psychological Review, 124(1), 91.

Fleming, S. M., & Lau, H. C. (2014). How to measure metacognition. Frontiers in Human Neuroscience, 8, 443. http://doi.org/10.3389/fnhum.2014.00443

Fleming, S. M., Huijgen, J., & Dolan, R. J. (2012). Prefrontal Contributions to Metacognition in Perceptual Decision Making. Journal of Neuroscience, 32(18), 6117–6125. http://doi.org/10.1523/JNEUROSCI.6489-11.2012

Fleming, S. M., Ryu, J., Golfinos, J. G., & Blackmon, K. E. (2014). Domain-specific impairment in metacognitive accuracy following anterior prefrontal lesions. Brain, 137(Pt 10), 2811–2822. http://doi.org/10.1093/brain/awu221

Fleming, S. M., Weil, R. S., Nagy, Z., Dolan, R. J., & Rees, G. (2010). Relating introspective accuracy to individual differences in brain structure. Science, 329(5998), 1541–1543. http://doi.org/10.1126/science.1191883

Galvin, S. J., Podd, J. V., Drga, V., & Whitmore, J. (2003). Type 2 tasks in the theory of signal detectability: discrimination between correct and incorrect decisions. Psychonomic Bulletin & Review, 10(4), 843–876.

Gelman, A., & Hill, J. (2007). Data Analysis Using Regression and Multilevel/Hierarchical Models. Cambridge University Press.

Gelman, A., & Rubin, D. B. (1992). Inference from iterative simulation using multiple sequences. Statistical Science, 7(4), 457–472. http://doi.org/10.1214/ss/1177011136

Goupil, L., Romand-Monnier, M., & Kouider, S. (2016). Infants ask for help when they know they don't know. Proceedings of the National Academy of Sciences, 113(13), 3492–3496. http://doi.org/10.1073/pnas.1515129113

Hautus, M. J. (1995). Corrections for extreme proportions and their biasing effects on estimated values of d'. Behavior Research Methods Instruments and Computers, 27, 46–46.

Heatherton, T. F. (2011). Neuroscience of self and self-regulation. Annual Review of Psychology, 62, 363–390. http://doi.org/10.1146/annurev.psych.121208.131616

Howell, D. C. (2009). Statistical methods for psychology. Wadsworth Pub Co.

Jang, Y., Wallsten, T. S., & Huber, D. E. (2012). A stochastic detection and retrieval model for the study of metacognition. Psychological Review, 119(1), 186.

Keene, O. N. (1995). The log transformation is special. Statistics in Medicine, 14(8), 811–819.

Kelemen, W. L., Frost, P. J., & Weaver, C. A. (2000). Individual differences in metacognition: evidence against a general metacognitive ability. Memory & Cognition, 28(1), 92–107.

Ko, Y., & Lau, H. (2012). A detection theoretic explanation of blindsight suggests a link between conscious perception and metacognition. Philosophical Transactions of the Royal Society B: Biological Sciences, 367(1594), 1401–1411.

Kruschke, J. K. (2014). Doing Bayesian Data Analysis. Academic Press.

Lau, H. C., & Rosenthal, D. (2011). Empirical support for higher-order theories of conscious awareness. Trends in Cognitive Sciences, 1–9. http://doi.org/10.1016/_j.tics.2011.05.009

Lee, M. D. (2008). BayesSDT: software for Bayesian inference with signal detection theory. Behavior Research Methods, 40(2), 450–456.

Lee, M. D., & Wagenmakers, E.-J. (2014). Bayesian Cognitive Modeling: A Practical Course. Cambridge University Press.

Lichtenstein, S., Fischhoff, B., & Phillips, L. D. (1982). Calibration of probabilities: The state of the art to 1980. In D. Kahneman, P. Slovic, & A. Tversky (Eds.), Judgment under uncertainty: Heuristics and biases. Cambridge University Press.

Macmillan, N., & Creelman, C. (2005). Detection theory: a user's guide. New York: Lawrence Erlbaum.

Maniscalco, B., & Lau, H. (2014). Signal Detection Theory Analysis of Type 1 and Type 2 Data: Meta-d“, Response-Specific Meta-d,” and the Unequal Variance SDT Model. In S. M. Fleming & C. D. Frith (Eds.), The Cognitive Neuroscience of Metacognition. Springer.

Maniscalco, B., & Lau, H. C. (2012). A signal detection theoretic approach for estimating metacognitive sensitivity from confidence ratings. Consciousness and Cognition, 21(1), 422–430. http://doi.org/10.1016/_j.concog.2011.09.021

Masson, M. E. J., & Rotello, C. M. (2009). Sources of bias in the Goodman-Kruskal gamma coefficient measure of association: implications for studies of metacognitive processes. Journal of Experimental Psychology. Learning, Memory, and Cognition, 35(2), 509–527. http://doi.org/10.1037/a0014876

Matzke, D., Lee, M.D., & Wagenmakers, E.-J. (2014) Signal detection theory: Parameter expansion. In M. D. Lee & E.-J. Wagenmakers (Eds.), Bayesian Cognitive Modeling: A Practical Course (pp. 187–195). Cambridge University Press.

McCurdy, L. Y., Maniscalco, B., Metcalfe, J., Liu, K. Y., de Lange, F. P., & Lau, H. C. (2013). Anatomical coupling between distinct metacognitive systems for memory and visual perception. Journal of Neuroscience, 33(5), 1897–1906. http://doi.org/10.1523/JNEUROSCI.1890-12.2013

Moeller, S. J., & Goldstein, R. Z. (2014). Impaired self-awareness in human addiction: deficient attribution of personal relevance. Trends in Cognitive Sciences, 0(0). http://doi.org/10.1016/j.tics.2014.09.003

Nelson, T. (1984). A comparison of current measures of the accuracy of feeling-of-knowing predictions. Psychological Bulletin, 95, 109–133.

Norton, E. H., Fleming, S. M., Daw, N. D., & Landy, M. S. (2017). Suboptimal criterion learning in static and dynamic environments. PLoS Computational Biology, 13(1), e1005304.

Palmer, E. C., David, A. S., & Fleming, S. M. (2014). Effects of age on metacognitive efficiency. Consciousness and Cognition, 28(1), 151–160. http://doi.org/10.1016/_j.concog.2014.06.007

Persaud, N., McLeod, P., & Cowey, A. (2007). Post-decision wagering objectively measures awareness. Nature neuroscience, 10(2), 257–261.

Plummer, M. (2003). JAGS: A program for analysis of Bayesian graphical models using Gibbs sampling. Presented at the Proceedings of the 3rd International Workshop on Distributed Statistical Computing.

Pouget, A., Drugowitsch, J., & Kepecs, A. (2016). Confidence and certainty: distinct probabilistic quantities for different goals. Nature Neuroscience, 19(3), 366–374. http://doi.org/10.1038/nn.4240

Rabbitt, P., & Vyas, S. (1981). Processing a display even after you make a response to it. how perceptual errors can be corrected. The Quarterly Journal of Experimental Psychology Section A, 33(3), 223–239. http://doi.org/10.1080/14640748108400790

Rahnev, D., Koizumi, A., McCurdy, L. Y., D’Esposito, M., & Lau, H. C. (2015). Confidence Leak in Perceptual Decision Making. Psychological Science, 0956797615595037. http://doi.org/10.1177/0956797615595037

Rausch, M. & Zehetleitner, M. (2016) Visibility is not equivalent to confidence in a low contrast orientation discrimination task. Frontiers in Psychology, 22(7), 591.

Schooler, J. W. (2002). Re-representing consciousness: Dissociations between experience and meta-consciousness. Trends in Cognitive Sciences, 6(8), 339–344.

Scott, R. B., Dienes, Z., Barrett, A. B., Bor, D., & Seth, A. K. (2014). Blind insight: metacognitive discrimination despite chance task performance. Psychological Science, 25(12), 2199–2208. http://doi.org/10.1177/0956797614553944

Spiegelhalter, D. J., Best, N. G., Carlin, B. P., & Van Der Linde, A. (2002). Bayesian measures of model complexity and fit. Journal of the Royal Statistical Society: Series B (Statistical Methodology), 64(4), 583–639. http://doi.org/10.1111/1467-9868.00353

Treisman, M. (1984). A theory of criterion setting: an alternative to the attention band and response ratio hypotheses in magnitude estimation and cross-modality matching. Journal of Experimental Psychology. General, 113(3), 443–463.

Weil, L. G., Fleming, S. M., Dumontheil, I., Kilford, E. J., Weil, R. S., Rees, G., et al. (2013). The development of metacognitive ability in adolescence. Consciousness and Cognition, 22(1), 264–271. http://doi.org/10.1016/j.concog.2013.01.004

Weiskrantz, L., Warrington, E. K., Sanders, M. D., & Marshall, J. (1974). Visual capacity in the hemianopic field following a restricted occipital ablation. Brain, 97(1), 709–728.

Wiecki, T. V., Sofer, I., & Frank, M. J. (2013). HDDM: Hierarchical Bayesian estimation of the Drift-Diffusion Model in Python. Frontiers in Neuroinformatics, 7, 14. http://doi.org/10.3389/fninf.2013.00014

